# VASC: dimension reduction and visualization of single cell RNA sequencing data by deep variational autoencoder

**DOI:** 10.1101/199315

**Authors:** Dongfang Wang, Jin Gu

## Abstract

Single cell RNA sequencing (scRNA-seq) is a powerful technique to analyze the transcriptomic heterogeneities in single cell level. It is an important step for studying cell sub-populations and lineages based on scRNA-seq data by finding an effective low-dimensional representation and visualization of the original data. The scRNA-seq data are much noiser than traditional bulk RNA-Seq: in the single cell level, the transcriptional fluctuations are much larger than the average of a cell population and the low amount of RNA transcripts will increase the rate of technical dropout events. In this study, we proposed VASC (deep Variational Autoencoder for scRNA-seq data), a deep multi-layer generative model, for the unsupervised dimension reduction and visualization of scRNA-seq data. It can explicitly model the dropout events and find the nonlinear hierarchical feature representations of the original data. Tested on twenty datasets, VASC shows superior performances in most cases and broader dataset compatibility compared with four state-of-the-art dimension reduction methods. Then, for a case study of pre-implantation embryos, VASC successfully re-establishes the cell dynamics and identifies several candidate marker genes associated with the early embryo development.

## Background

Characterizing the cellular states in single cell level is crucial for understanding the cell-cell heterogeneities and the biological mechanisms not observed by the average behaviors of a bulk of cells. Single cell RNA sequencing (scRNA-seq) is a promising high-throughput technique to simultaneously profile the transcriptomes of a large number of individual cells [1]. Thousands of genes are expressed in a single cell at the same time. Their expression levels are usually tightly regulated regarding to a limited number of cellular states. Finding an effective low-dimensional representation of the scRNA-seq data is the basic step for the data visualization and the downstream analysis, such as the cell lineage establishment and the cell sub-population identification [2]. Currently, several traditional dimension reduction methods used for the bulk RNA-Seq data, such as PCA [3] and t-SNE [4], are still widely used for the scRNA-seq data. However, the transcriptional burst effects and the low amount of RNA transcripts in single cells make the scRNA-seq data much noisier than the bulk RNA-Seq. For example, the scRNA-seq data have many unexpected dropout events (many data points are zero or near-zero) [5]. These noises make those traditional methods work inefficiently. To improve the analysis, one useful strategy is to explicitly mimic the data generation process by a probabilistic model. For example, the zero-inflated factor analysis (ZIFA) [6], which combines the probabilistic factor analysis with conditional dropout probability, is developed to find the latent low dimension subspace. However, ZIFA can only model linear patterns by a single hidden layer, which limits its performances on the datasets with complex cellular states in the original data space. Another strategy is to embed the cells into another low-dimensional space by preserving the cell-cell similarity (or distance) in the original data space. But, this kind of methods, such as SIMLR [7], frequently change the basic topological information in the embedded space.

In recent years, deep probabilistic hidden models show superior performances for representing complex features of high dimensional data, especially for images and speeches.[8, 9] In this study, we developed a deep model VASC (deep variational autoencoder for scRNA-seq data) to analyze and visualize scRNA-seq data. It can capture non-linear variations and automatically learn a hierarchical representation of the input data. And, it uses the Gumbel distribution to better model the zero and near-zero dropout events. We systematically compared VASC with several state-of-the-art dimension reduction methods on twenty datasets. Results show that VASC has superior performances in most cases and broader dataset compatibility.

## Results

### VASC: the method overview

VASC, a deep variational autoencoder [9-11] based generative model, was designed to find an effective low-dimensional representation and facilitate the visualization of scRNA-seq datasets. It modeled the distribution of high-dimensional original data P(***X***), by a set of latent variables ***z*** (the dimension of ***z*** should be much smaller than ***X***, in particular, as few as two dimensions for the visualization). Its objective was to find the optimal ***z*** in terms of capturing the intrinsic information of the input data. In a probabilistic view, the posterior distribution P(***z*** |***X***) could be treated as the best distribution of ***z*** with the observed data ***X***. However, P(***z*** |***X***) is usually intractable. Variational inference is proposed to solve this problem by designing another common distribution family Q(***z*** |***X***) (called as variational distribution) to approximate P(***z*** |***X***). The minimization of the Kullback–Leibler (KL) divergence between the two distributions is usually used for the approximation. The variational distribution Q(***z*** |***X***) should have enough representation capacity to model the complex information of P(***z*** |***X***) in the scRNA-seq dataset, and on the other hand, should be tractable for efficient computation. In VASC, deep neural networks were used to explicitly model the variational distribution Q(***z*** |***X***). Unlike the traditional varitional inference methods, deep neural networks can approximate arbitrary functions and can be solved efficiently with the stochastic gradient descent methods.

Generally, VASC had three major parts, called the encoder network, the decoder network and the zero-inflated layer (Figure 1). The encoder network, designed as a three-layer neural network, generated the parameters of the variational distribution. Note that before the first layer, we added a “dropout” noise layer [12], which randomly set some data points as zero in the original expression matrix. This layer can be treated as an artificial mimic of the “dropout” events – the expressions of some genes are dropped to zero. From a computational view, it introduced additional random noises for the sample training, which can reduce the overfitting risk during the learning process. We supposed a multi-dimensional Gaussian distribution Q(***z*** |***X***) of latent variables ***z*** given the expression values ***X***, whose mean and variance parameters could be generated by the encoder network. Then, the learned Q(***z*** |***X***) was used to re-generate pseudo samples ***X***’ by the decoder network, another three-layer neural network. Finally, a zero-inflation layer, based on a double-exponential distribution, was designed to mimic the dropout events by randomly setting some data points as zero [6, 13]. The Gumbel distribution was used in this layer instead of the conditional binomial distribution for the back-propagation [14-16]. VASC was optimized by a stochastic gradient descent-based RMSprop methods [17]. The objective was to minimize an auxiliary loss function of the K-L divergence between Q(***z*** |***X***) and P(***z*** |***X***). After the auto-encoding procedure, a 2D representation was learned for the visualization and the downstream analysis.

**Figure 1.**
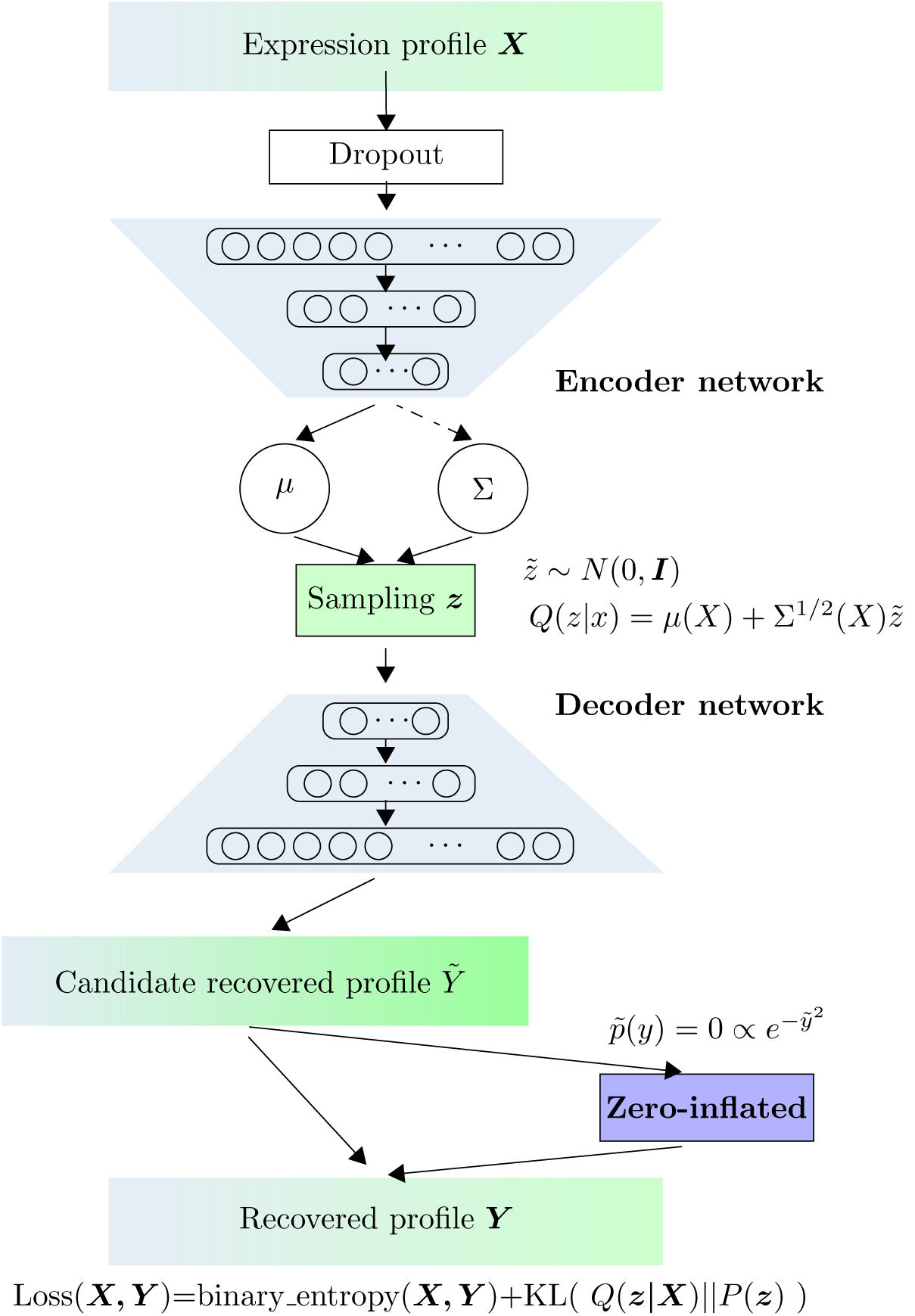
Overview of VASC method. VASC consists of three parts: the encoder network, the decoder network and the zero-inflated layer. Both the encoder and decoder networks are designed as three-layers fully connected neural networks.

### Visualization and performance comparison

We tested VASC and four state-of-the-art dimension reduction methods (PCA [3], t-SNE [4], ZIFA [6] and SIMLR [7]) on twenty datasets with different number of cells and sequencing protocols. Firstly, we compared the 2D visualizations on six “golden” datasets (these datasets provide high-confident cell labels), which had the number of cells range from tens to more than one thousand (Figure 2). The top three datasets in the figure 2, Goolam [18], Biase [19] and Yan [20], studied the embryonic development from zygote cells to blast cells. The three methods PCA, ZIFA and VASC could roughly re-establish the lineages of different cell types. (The cells were expected to be arranged by the order of zygote, 2-cell, 4-cell, 8-cell, 16-cell and blast cells) But, t-SNE and SIMLR, both of which use neighbor preserving embedding, showed poor results on these datasets. For the Goolam dataset, VASC further separated 16-cell and blast with 8-cell stages. For the Biase dataset, VASC better separated blast cells with 4-cell stages, and identified one zygote as a possible outlier. For the Yan dataset, VASC better separated 4-cell from zygote and 2-cell stages. These results showed that VASC can better model the cell lineages during embryo development than PCA and ZIFA.

**Figure 2.**
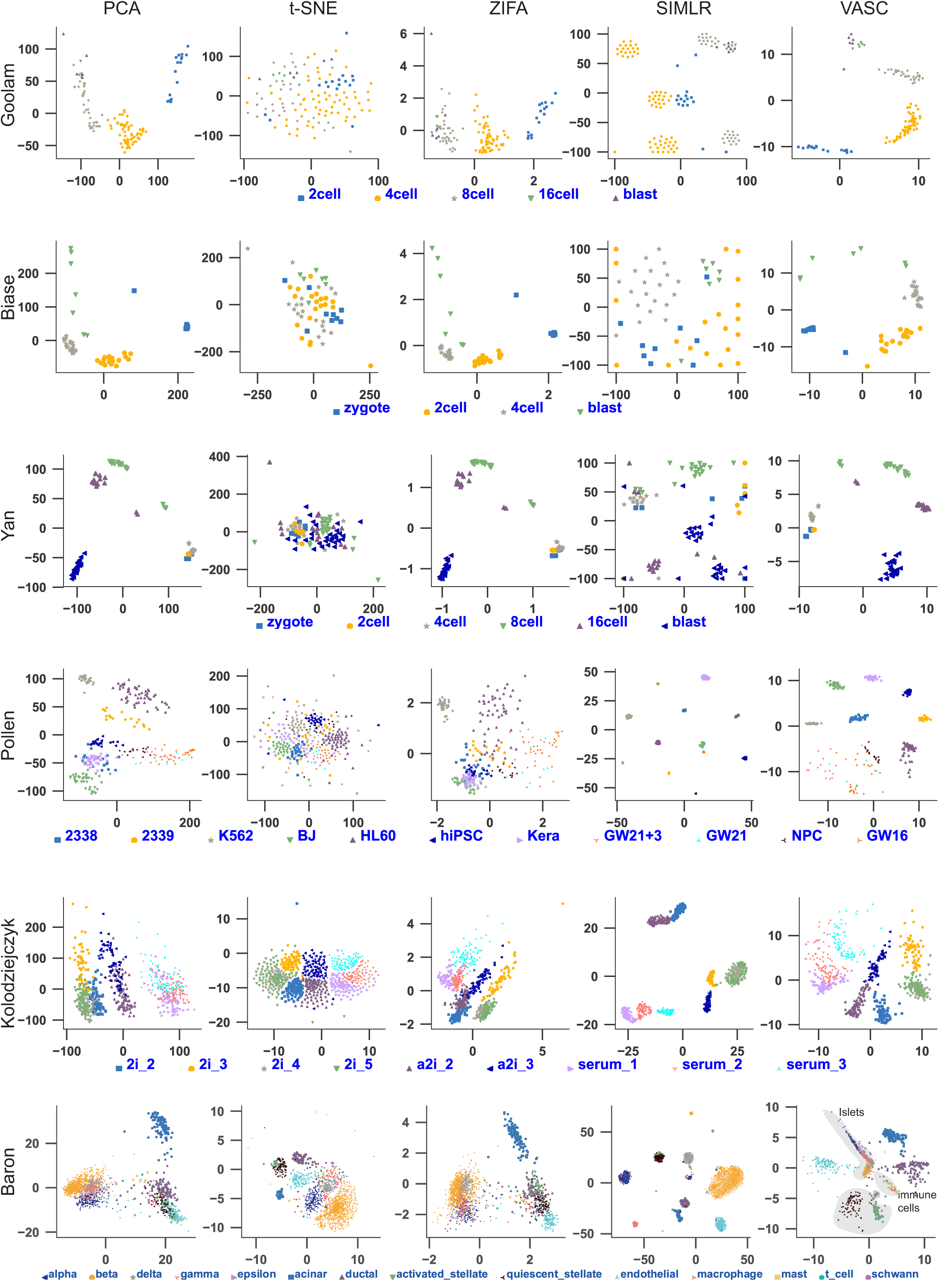
Visualization of scRNA-seq datasets by different methods. Each point represents a cell. Different cell types are marked by different colors and shapes. The visualizations of the other datasets are provided in supplementary materials.

The fourth dataset, Pollen [21], sequenced eleven different cell types. In this case, PCA and ZIFA showed inferior performances. In the SIMLR visualization, eleven compact clusters of cells were formed, but at least four clusters were composed of more than one cell types. (The points of different cell types were stacked together for possible misleading visualization) This result was undesirable because the cells from different types should not compactly cluster together. Instead, VASC formed eight compact clusters of cells from the same cell type. The other three cell types, GW16, GW21 and GW21+3 (originally sampled from the germinal zone of human cortex at gestational week 16, 21 and cultured for another three weeks respectively), were distributed more decentralized than the others. These cells, along with NPCs (neural progenitor cells), are all neural cells, and were more closely represented by VASC, which should be more reasonable.

The Kolodziejczyk dataset [22] studied embryonic stem cells grown under three different conditions: serum, 2i, and alternative 2i (a2i). However, for every condition, there existed different experimental batches. The visualizations showed that t-SNE and VASC had better results: PCA separated the cells grown in the three different conditions but almost mixed the batches; ZIFA better separated the cells in different growth conditions and batches but incorrectly mixed one 2i cell batch (2i_2) with a2i cells; SIMLR separated most cell populations with different growth conditions and batches (except two batches of 2i cells) but incorrectly grouped the cells from 2i and a2i conditions; and, only t-SNE and VASC can separated most cell populations and maintained their relative correct positions at the same time.

The Baron dataset [23] included several sequencing subsets from four human donors and two mice, and the figure 1 showed the results of the first donor with 1,937 cells from 14 different cell types. On this dataset, PCA and ZIFA only separated few cell types. t-SNE and SIMLR got similar separation results, although SIMLR produced more compact clusters. Putative clusters of SIMLR contained different extents of mixtures of different cell types. (For example, two kinds of stellate cells were completely mixed) From the 2D visualization, obviously, VASC showed better separation of different cell types. And the cells from close cell lineages were clustered together: the alpha, beta, delta, gamma and epsilon cells, all within islets, were close to each other; the beta cells with the largest number (872 cells) were most compactly clustered by VASC; three types of immune cells, macrophage (14 cells), mast (8 cells) and t_cells (2 cells), were close to each other; and the schwann cells with only 5 samples were almost separated (please see the purple dots in the central region).

Then, the performances of these methods were quantitatively assessed by comparing the cell sub-populations in the reduced subspaces (the sub-populations were identified by *k*-means clustering [24]) with the true cell type labels annotated by the original references. Four different indexes – NMI (normalized mutual information score) [25], ARI (adjusted rand index) [26], HOM (homogeneity) and COM (completeness) [27] were used to quantitatively assess the clustering performances. (Please see the Methods section for details) These comparisons showed that VASC outperformed the compared methods in terms of NMIs and ARIs in most cases (best performances on 15 and 17 datasets, respectively) (Figure 3). And, VASC always ranked in the top two methods on all the tested datasets, which suggested that it has broad compatibility for various kinds of scRNA-seq datasets. Please see the detailed results in the supplementary materials.

**Figure 3.**
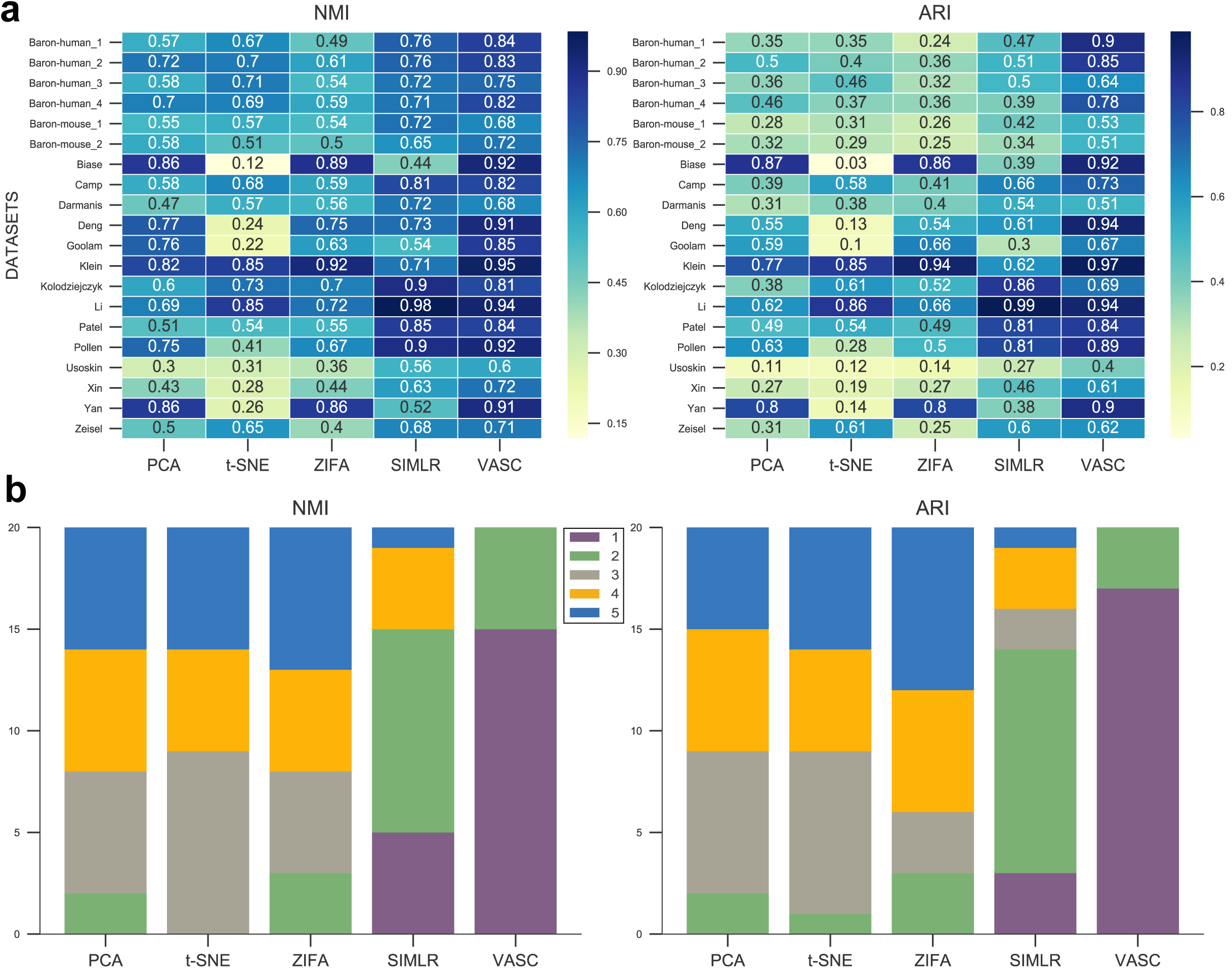
Performance comparison of different methods. (a) The NMIs and ARIs for each method on each dataset. (b) The statistics of the ranks of the compared methods based on NMIs and ARIs.

### Analysis of the model stability and parameter setting

In this section, we analyzed the stability and several parameter settings of VASC. Firstly, we analyzed the model fitting processes of VASC on two datasets, the Pollen and Biase datasets (with 301 and 56 cells, respectively). The loss function decreased sharply during the first a few epochs, and simultaneously, NMI/ARI values increased sharply (Figures 4a & 4b). After the first 100 epochs, the loss curves quickly converged to a lower limit and the loss fluctuations of the dataset with more samples (Pollen) were smaller than the one with fewer samples (Biase). Based on these observations, the stopping criterion for VASC was set as that the loss function had no obvious decrease within 50 epochs (see details in Methods).

**Figure 4.**
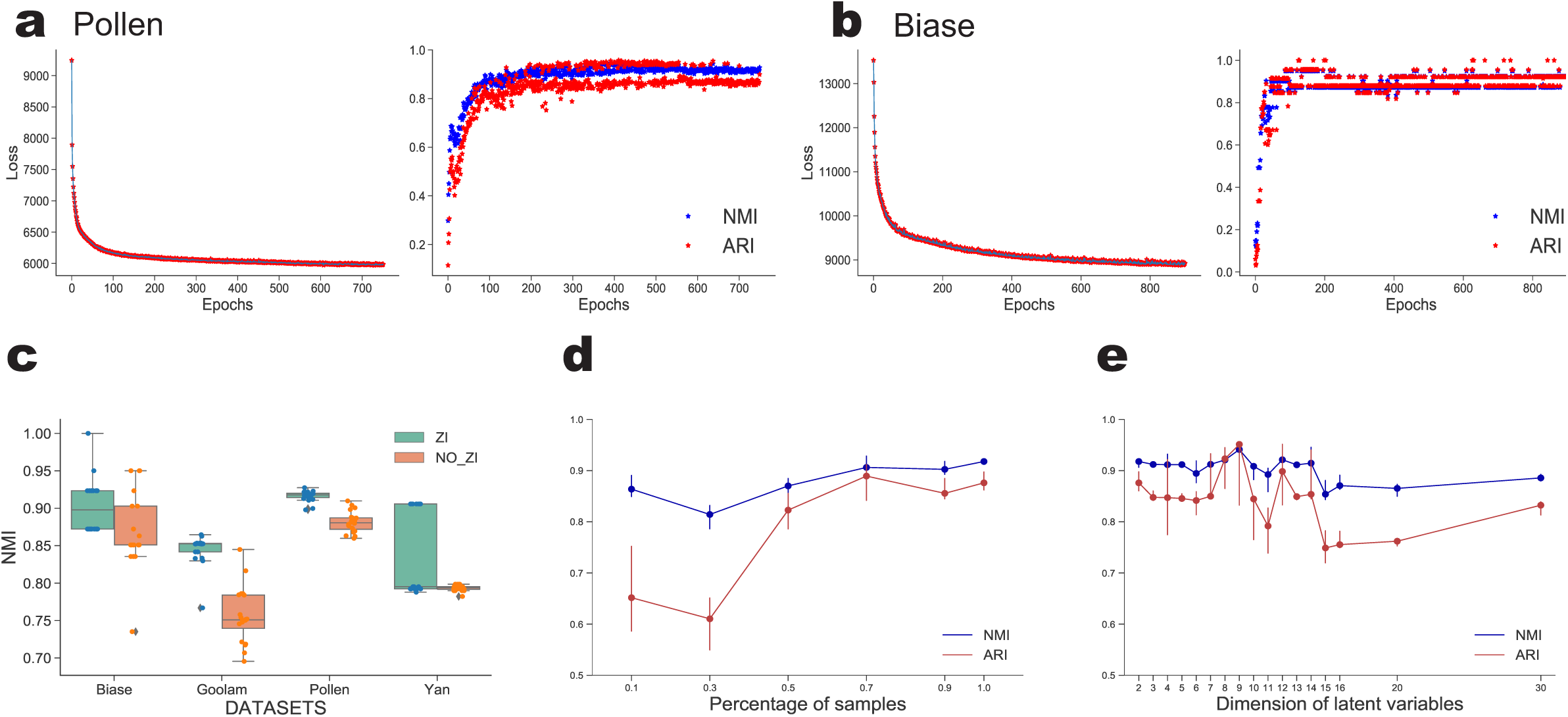
Analysis of VASC algorithm. (a) The iteration process of the Pollen dataset. The left part is the change of loss values versus epochs and the right part is the change of NMI and ARI versus epochs. (b) The iteration process of the Biase dataset. (c) The stability of VASC. The boxplots were based 20 repeated runs with and without the zero-inflation layer. (d) The down-sampling experiments based on the Pollen dataset. (e) The effects of the dimensions (from 2 to 30) for the latent variables based on the Pollen dataset.

Next, the stability of VASC was analyzed. Due to the randomness of the stochastic gradient descent method, the model initialization and the of *k*-means clustering, different runs generated slight different results. The four datasets with the fewest sample sizes, Biase (56 samples), Goolam (124), Pollen (301) and Yan (90) were used to test the stability of VASC by 20 repeated runs. As expected, the two dataset sets with more samples (Goolam & Pollen) showed much higher consistent results (Figure 4c). The following down-sampling analysis of the Pollen dataset also suggested the same trend (Figure 4d). The NMIs of the Biase dataset were almost distributed between two values. Considering the relatively small number of cells (only 56 samples), this distribution may be caused by the turnover of just one or two cells at the boundary. The similar result was also observed for the Yan dataset. However, the Goolam and Pollen datasets with more cells did not show this pattern. Then, the down-sampling experiment based on the Pollen dataset was implemented to further test the method’s stability. The dataset was bootstrapped with 10%, 30%, 50%, 70%, 90% and 100% cells also with 20 repeated runs. It is observed that the NMIs and ARIs dropped if the number of samples were too small and their variations increased accordingly. But, VASC get comparable results if the percentage of sampled cells was above 50%.

Then, the effectiveness of the ZI layer, which modeled the dropout event, was assessed. Results showed that it improved both the stability and the average performances (Figure 4c).

The data projection to a 2D subspace is suitable for the visualization, but the subspace with higher dimension may explain more variations. In the model, the dimensions of the latent variables were changed from 2 to 20 for the Pollen dataset. Results showed that the increase of the dimensions did not improve the identification of known cell populations and the subspaces with too high dimensions even caused worse results (Figure 4e).

### Case study: human pre-implantation embryos

The scRNA-seq is very useful for studying the cell dynamics during embryo pre-implantation development. We applied VASC on a recently published dataset of human pre-implantation embryos (the Petropoulus dataset), including 1,529 cells with detailed annotations of developmental stages, inferred lineage and inferred pseudo-time information [28]. The 2D visualizations showed that VASC and t-SNE recovered the known developmental stages (form E3 to E7) more precisely, with the exception that the E3 cells were out of the trajectory by t-SNE. Both PCA and ZIFA generally recovered the stage trajectory but the E6 and E7 cells were largely overlapped. SIMLR, which emphasized the modularity of cell populations, did not re-establish the basic shapes. (Figures 5a-e)

**Figure 5.**
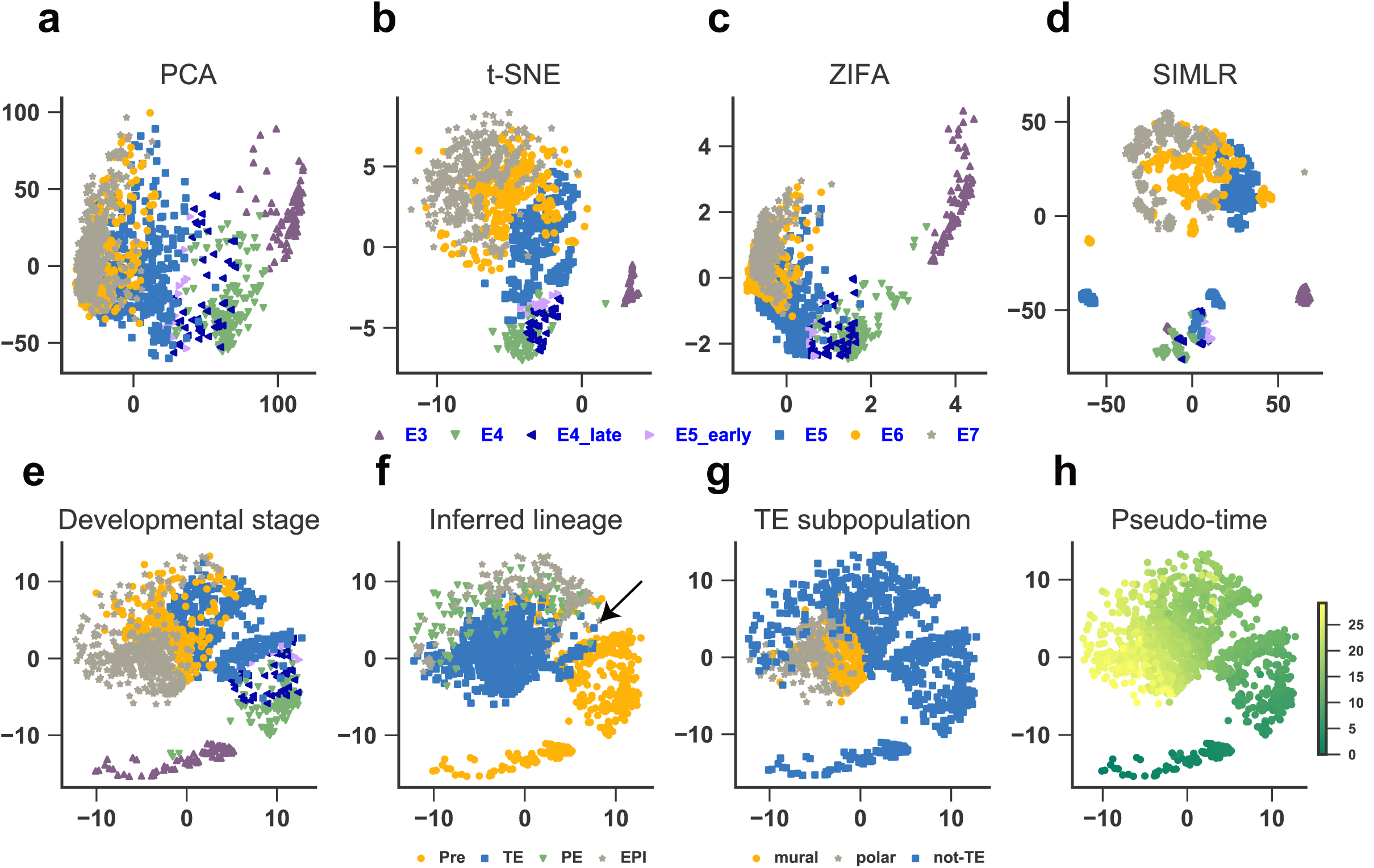
Case study: Petropoulos dataset. (a-e) The 2-D visualizations of PCA, t-SNE, ZIFA, SIMLR and VASC. The cells are annotated with the developmental stages. (f) The cells are annotated as pre-lineage and the other cells. (g) The TE cells are further annotated as mural and polar cells. (h) The cells are annotated with the inferred pseudo time.

Next, we investigated other annotations based on the visualizations. A sharper split was found in the E5 cells by VASC, compared with the t-SNE result. We re-annotated the cells with their lineages instead of the developmental stages. It is found that the sharp split learned by VASC was a good separation of the pre-lineage cells from the others (Figure 5f). The inner cell mass (ICM), including the PE (primitive endoderm) and EPI (epiblast), were split from TE (trophectoderm), and the boundary was almost perpendicular to the direction of developmental stage (Figure 5f). It was also found the two sub-populations of the TE cells, mural and polar cells, can be separated in the visualization (Figure 5g). Finally, the trajectory recovered by VASC was strongly coincided with the inferred pseudo time (Figure 5h).

The candidate genes associated with the embryo pre-implantation development were identified by calculating the Spearman’s correlations between the gene expressions and the two features in the reduced subspace. Many known regulators and markers were found in the top correlated genes, such as PGF, GCM1, CYP19A1, MUC15, CD24, CCR7, GREM2, CGA, GATA2, TDGF1, ESRG, GDF3 and DNMT3L mentioned in the dataset article (rank <= 100 for either feature). Interestingly, the top-ranked genes were significantly enriched in metabolic processes, such as “carbohydrate derivative metabolic process” (37 genes, q-value 5.63e-05 by DAVID 6.8 [29]), “oxidation-reduction process” (32 genes, 4.87e-05) and “lipid metabolic process” (32 genes, q-value 4.94e-03). Recent technique advances found several metabolic pathways play as essential factors for regulating stem cells’ stemness and differentiation [30]. These analyses identified several interesting candidate genes involved in different metabolic processes: for example, CYP11A1 (a member of the cytochrome P450 superfamily of enzymes, the sample superfamily of CYP19A1), NR2F2 (a member of the steroid thyroid hormone superfamily of nuclear receptors), PKM (a pyruvate kinase that catalyzes the transfer of a phosphoryl group from phosphoenolpyruvate to ADP, generating ATP and pyruvate, a key kinase for glycolysis), PPARG (a member of the peroxisome proliferator-activated receptor subfamily of nuclear receptors) and IDH1 (an isocitrate dehydrogenase, key enzyme for cytoplasmic NADPH production).

## Discussion

Dimension reduction (or low-dimensional representation) is the basic step for the visualization and the downstream analysis of scRNA-seq data. The four compared methods and VASC could be broadly divided as two categories: PCA, ZIFA and VASC aim at finding the representation which can best explain the variations of the original data, while t-SNE and SIMLR try to find another embedded space which can preserve the neighborhood relationship of the samples in the original space. According to our experiments, the former three methods can better keep the basic shapes of the data distributions. ZIFA can be treated as a combination of probabilistic PCA and the zero-inflated model. The major limitation is that it assumes a linear relationship between the hidden subspace and the observed data. VASC can deal with complex non-linear patterns based on deep neural networks. Results show that VASC has better performances than PCA and ZIFA, especially when the sample sizes are larger. The two embedding methods, t-SNE and SIMLR, frequently change the topology of the original data space. t-SNE tends to “disperse” the cells in the embedded subspace. Compared with t-SNE, SIMLR adds penalties on the modularity of samples in the embedded subspace, which forces the diagonal-block structure of the learned cell-cell similarity matrix, and tends to generate compact clusters. This penalty is very useful to identify the cell populations with distinct transcriptomes (for examples, the Pollen dataset). Nevertheless, it frequently fails, if the dataset studies “continuous” cell developmental processes or cell lineages. Overall, our experiments show that VASC is superior in most cases and broader dataset compatibility.

One major application of scRNA-seq is to analyze different cell types in single cell level. According to the quantitative results shown in the figure 3 two dimensions are enough to capture the major differences between different cells in most cases (NMI > 0.7 for 16 of 20 datasets by VASC). Although higher dimensions can explain more variations in the original datasets, additional variations not associated cell type information (for example, the fluctuations associated with cell cycle) may even reduce the separation of different cell types according to our experiments. The determination of the optimal dimension is a tricky task if the prior knowledge is limited. Usually, higher dimensions should be used if we want to study more subtle differences, for example, the intra-cell-type heterogeneity.

There are two parameters (the mean vector and the co-variance matrix) in the variational distribution Q(***z*** |***X***). When the sample size is small, it is better to fix the co-variance matrix. But, when the size is large enough (above 1,000 according to our preliminary experiments), a co-variance matrix learnt from the data can get better results. It is expected that more complex variational distribution families can be tested in near future, as the sample size of scRNA-seq dataset is quickly increasing.

It is found that the inclusion of zero-inflation (ZI) layer improves the representation of VASC in terms of recovering the known cell types. Compared with ZIFA, the Gumbel distribution used by the ZI layer didn’t generate strictly zeroes, which may additionally model the near-zero dropout events, which is proposed as a limitation of ZIFA [6].

The stochastic optimization algorithms used for the VASC model learning introduces variations in the dimension reduction results. Repeated runs are recommended for more consensus results, although this random effect is small if the sample size is over several hundreds. The running time is a common issue for deep models. For the large dataset with several thousands of cells, it cost several hours for the VASC model learning by a desktop-level computer with single GPU card, which should be acceptable for most scRNA-seq studies.

## Conclusions

In this study, a dimension reduction method VASC was developed for scRNA-seq data visualization and analysis. We systematically compared VASC with four state-of-the-art dimension reduction methods on twenty datasets. Results show that VASC achieves superior performances in most cases and is broadly suitable for different datasets with different data structures in the original space. The application on a dataset of the human pre-implantation embryo development shows that VASC can re-establish the cell dynamics in the reduced 2D-subspace and identify the associated marker genes.

## Methods

### Datasets

To demonstrate the performance of VASC, we analyzed twenty scRNA-seq datasets. All datasets were obtained via the website https://hemberg-lab.github.io/scRNA.seq.datasets/. They used ‘scater’ toolkit [31] for quality control. The human pre-implantation embryo data set (Petropoulus) with detailed annotations is obtained via ArrayExpress: E-MTAB-3929.

### Variational autoencoder

VASC is a deep variational autoencoder(VAE) based generative model and is designed for the visualization and low-dimensional representation analysis of scRNA-seq data. The variational autoencoder (VAE) aims to model the distribution P(***X***) of data points in a high-dimensional space, with the aid of low dimensional latent variables ***z***. The whole model could be divided into two procedures, the first of which is to generate the samples of ***z*** in the latent low-dimensional subspace, and the second is to map them to the original space. The critical point is to generate ***z*** having the high probability to recover the observed data matrix ***X***. It means that the generated ***z*** may possibly capture the intrinsic information of the original data. Theoretically, the best choice to generate ***z*** is the posterior distribution P(***z|X***), which however, is usually too complicated and intractable. VAE tries to use a varitional probability Q(***z|X***) to approximate it, which is optimized to minimize the Kullback–Leibler(KL) divergence between Q(***z|X***) and P(***z|X***):

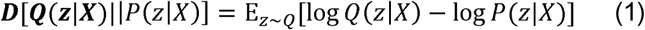

By applying the Bayes rule and rearranging the order, it can be re-written as:

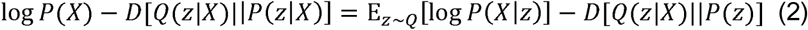

where P(***X***) is a constant and so to minimize the K-L divergence is equivalent to maximize the right-hand of the above equation. The right-hand has a natural autoencoder structure, with the encoder Q(***z|X***) from ***X*** to ***z*** and the decoder P(***X|z***) from ***z*** to ***X*** respectively. Two deep fully connected neural networks can be used to model these two parts.

### The VASC method

The whole VASC structure is shown in the figure 1. The model designs and the learning algorithms are described in details as below.

#### Input layer

VASC used the expression matrix from scRNA-seq data as inputs. As in all our experiments, no any gene filter was applied, and the whole expression matrix of the transciptome was directly fed to the model. This “no-filtering” pre-processing is a great help for the RNA-seq data analysis. The data were log-transformed to make the results more robust. The most important transformation, however, was to re-scale the expression of every gene in any single cell to [0,1] by dividing the maximum expression value of its own cell. Some subsequent designs were based on this pre-processing step.

#### Dropout layer

A dropout layer [12] was added immediately after the input layer. The dropout rate was set to 0.5, larger than the usual choice of deep models. This layer set some features as zeros during the encoding phase, which can increase the performance for the modeling learning [32]. This layer should be a good choice for scRNA-seq data because it may be regarded as artificial and additional “dropout” events.

#### Encoder network

The encoder network was designed as a three-layer fully connected neural network with decreasing dimensions 512,128 and 32. The first layer did not use non-linear activation, which acted as an embedded PCA transformation. Many complex algorithms, including t-SNE, get benefits from the PCA transformation. A L1-norm regularization was added for the weights in this layer, which penalized the sparsity of the model. The next two layers were accompanied by ReLU activation, which made the output sparse and stable for deep models [33].

#### Latent sampling layer

Latent variables ***z*** were modeled by a Gaussian distribution, with the standard normal prior N(0,***I***). The encoder network was used to estimate its posterior parameters. Usually, both the two parameters *μ*, **Σ** were needed to be estimated. A linear activation was used for the parameter *μ* estimation. According to our experiments, it was found that it was better to fix the **Σ** parameter and set log **Σ** = 1 if the dataset only has small sample size. For the datasets with large sample sizes (more than 1,000 cells), this parameter can also be trained by the encoder network. A ‘softplus’ activation was used for the estimation of log **Σ**. Because the neural network cannot have a stochastic layer, which could not be tackled by back-propagation algorithm. A re-parameterization trick was used to remove the randomness to inputs. It is easy to see, drawing a sample 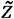 from *N*(*μ*, **Σ**) is equivalent to draw a sample from *N*(0,*I*) and then let 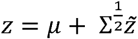 (Please see supplementary materials for more details)

#### Decoder network

The decoder network used the generated ***z*** to recover the original expression matrix, and was designed as a three-layer fully connected neural network with dimensions of hidden units 32, 128, and 512, and an output layer. The first three layers used ‘ReLU’ activations and the final layer with sigmoid to make the output within [0,1] (this is why the previous [0,1] re-scaling transformation must be applied).

#### Zero-inflation (ZI) layer

An additional ZI layer was added after the decoder network. Adapted from the model used by ZIFA [6], we modeled the dropout events by the probability 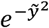 where 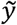 is the recovered expression value by the decoder network. Back-propagation, as said before, cannot deal with stochastic units, and meanwhile, it cannot deal with discrete units either. A Gumbel-softmax distribution [16] was introduced deal with these difficulties. Supposed the drop-probability *p, q*, = 1 – *p*, the sample from Gumbel-softmax distribution was obtained by:

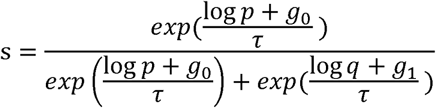

where g_0_, *g*_1_were sampled from a Gumbel(0,1) distribution. The samples could then be obtained by first drawing u ∼ Uniform (0,1) and then computing g = –log(– log *u*). As the hyper-parameter *τ* → 0, the generated samples from the Gumbel-softmax distribution should be identical to the samples from the Bernoulli distribution. In practice, too small *τ* makes the gradient too small and the optimization algorithm cannot work. Our experiments showed that the it would be better by setting *τ* between 0.5∼1 for the datasets of small sample size. For the datasets with more cells, an annealing strategy may get better results. (See supplementary materials for details)

#### Loss function

The loss function as shown in the equation (2), is composed of two components. The first part, because of the [0,1] scale of our data, was computed by binary cross-entropy loss function. The second part, controlling the divergence between posterior distribution and the prior *N*(0, *I*), could be computed analytically. (See supplementary materials for more details)

#### Optimization

The whole structure, now, could be optimized end-to-end using the stochastic gradient descent-based optimization algorithm. The RMSprop method [17] was chosen by VASC. And the learning rate was set as 0.0001, which ensured the convergence on all the tested datasets. The training processes was stopped if the training loss did not show obvious decrease within 50 epochs.

### Performance assessment

To measure the quality of low-dimensional representation, *k*-means clustering was applied to the 2D representations of all the methods and compared the clustering results with known cell types provided by their original references. The number of clusters, *k*, was set to number of known cell types. Four measure indices were used to assess the performances.

#### Normalized mutual information (NMI).[25]

Suppose P is the predicted clustering results, and T is the known cell types (the same below). Denote the entropy of P and T as H(P) and H(T), respectively, and the mutual information between them as MI(P,T). NMI is computed as:

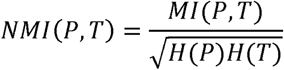

#### Adjusted rand index (ARI).[26]

Suppose *n* is the total number of samples, *a*_*i*_ is the number of samples appearing in the i-th cluster of P, *b*_*j*_ is the number of samples appearing in the j-th types of T, and *n*_*ij*_ is the number of overlaps between the i-th cluster of P and the j-th type and T. API is computed as:

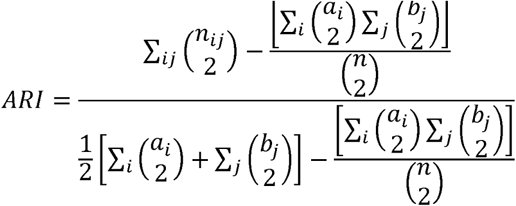

#### Homogeneity.[27]

This measure expects every cluster only contains samples from one cell type. Suppose H(T|P) is the cross-entropy of cell types given the cluster P. The Homogeneity score is computed by:

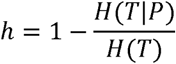

#### Completeness.[27]

This measure expects samples from one cell type are assigned to the same cluster, and is computed as:

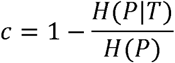

For all the above measures, larger values (up to 1) mean better performances.

### Benchmarking

For each dataset, we considered four state-of-the-art dimension reduction methods – PCA[3], t-SNE[4], ZIFA[6] and SIMLR[7]. For all the methods, no gene filtering was used and the same log-2 transformation was applied. For PCA and t-SNE, we used the built-in python sklearn package functions. For the datasets more than 500 cells, we firstly applied a PCA transformation with 500 dimensions before t-SNE. The key parameter of t-SNE – perplexity was set to 0.2 times the number of cells as the paper [28] suggested. For ZIFA, we downloaded their package and used their block_ZIFA module because of the large number of genes. For SIMLR, we used their R package. For benchmarking the dimension reduction performance, *k*-means was used to get the predicted cell types based on their 2D representations.

**Table 1.**
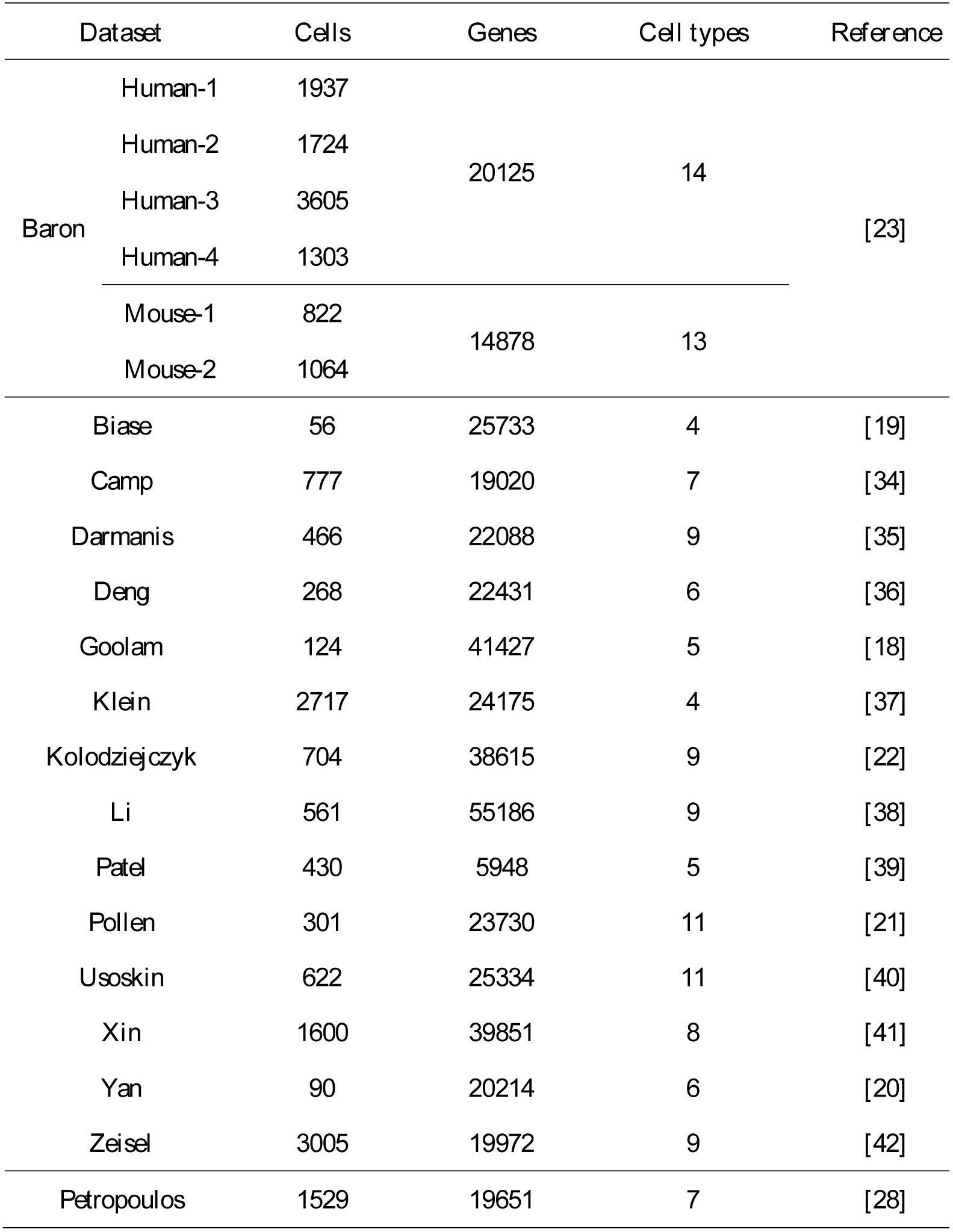
The list of scRNA-seq datasets used in this study

## Additional files

Supplementary materials include more details about VASC and supplementary figures.

### Abbreviations

PCA: principal components analysis
SIMLR: single-cell interpretation via multiple kernel learning.
t-SNE: t-distributed stochastic neighbor embedding
VAE: variational autoencoder
VASC: deep variational autoencoder for scRNA-seq
ZIFA: zero-inflated factor analysis

## Acknowledgements

We thank Xiangyu Li, Jianyang Zeng, Michael Zhang and Jun Li for their helpful discussions. We address special thanks to the share of single cell datasets by Hemberg group from the Wellcome Trust Sanger Institute.

## Availability of data and materials

All datasets used in this study are publicly available as stated in ‘Datasets’ section of ‘Methods’. Source codes implemented by keras (https://github.com/fchollet/keras) could be found via: https://github.com/wang-research/VASC.

## Authors’ contributions

DW and JG designed this study and developed the algorithm. DW made the detailed implementation and performed the data analysis. DW and JG wrote this manuscript. All authors read and approved the manuscript.

## Ethics approval and consent to participate

Not applicable.

## References

1. Shapiro, E., T. Biezuner, and S. Linnarsson, Single-cell sequencing-based technologies will revolutionize whole-organism science. Nature reviews. Genetics, 2013. 14(9): p., 618.

2. Stegle, O., S.A. Teichmann, and J.C. Marioni, Computational and analytical challenges in single-cell transcriptomics. Nature reviews. Genetics, 2015. 16(3): p., 133.

3. Wold, S., K. Esbensen, and P. Geladi, Principal component analysis. Chemometrics and intelligent laboratory systems, 1987. 2(1-3): p. 37–52.

4. Maaten, L.v.d. and G. Hinton, Visualizing data using t-SNE. Journal of Machine Learning Research, 2008. 9(Nov): p. 2579–2605.

5. Bacher, R. and C. Kendziorski, Design and computational analysis of single-cell RNA-sequencing experiments. Genome biology, 2016. 17(1): p., 63.

6. Pierson E. and C. Yau, ZIFA: Dimensionality reduction for zero-inflated single-cell gene expression analysis. Genome biology, 2015. 16(1): p., 241.

7. Wang, B., et al., SIMLR: a tool for large-scale single-cell analysis by multi-kernel learning. arXiv preprint arXiv:1703.07844, 2017.

8. Hinton, G.E. and R.R. Salakhutdinov, Reducing the dimensionality of data with neural networks. science, 2006. 313(5786): p. 504–507.

9. Kingma, D.P. and M. Welling, Auto-encoding variational bayes. arXiv preprint arXiv:1312.6114, 2013.

10. Kingma, D. and M. Welling. Efficient gradient-based inference through transformations between bayes nets and neural nets. in International Conference on Machine Learning., 2014.

11. Doersch, C., Tutorial on variational autoencoders. arXiv preprint arXiv:1606.05908, 2016.

12. Srivastava, N., et al., Dropout: a simple way to prevent neural networks from overfitting. Journal of Machine Learning Research, 2014. 15(1): p. 1929–1958.

13. Kharchenko P.V., L. Silberstein, and D.T. Scadden, Bayesian approach to single-cell differential expression analysis. Nature methods, 2014. 11(7): p. 740–742.

14. Gumbel, E.J., Statistical theory of extreme values and some practical applications: a series of lectures. 1954: US Govt. Print. Office.

15. Maddison, C.J., D. Tarlow, and T. Minka. A* sampling. in Advances in Neural Information Processing Systems., 2014.

16. Jang, E., S. Gu, and B. Poole, Categorical reparameterization with gumbel-softmax. arXiv preprint arXiv:1611.01144, 2016.

17. Tieleman, T. and G. Hinton, Lecture, 6.5-rmsprop: Divide the gradient by a running average of its recent magnitude. COURSERA: Neural networks for machine learning, 2012. 4(2): p. 26–31.

18. Goolam, M., et al., Heterogeneity in Oct4 and Sox2 targets biases cell fate in 4-cell mouse embryos. Cell, 2016. 165(1): p. 61–74.

19. Biase, F.H., X. Cao, and S. Zhong, Cell fate inclination within 2-cell and 4-cell mouse embryos revealed by single-cell RNA sequencing. Genome research, 2014. 24(11): p. 1787–1796.

20. Yan, L., et al., Single-cell RNA-Seq profiling of human preimplantation embryos and embryonic stem cells. Nature structural & molecular biology, 2013. 20(9): p. 1131–1139.

21. Pollen, A.A., et al., Low-coverage single-cell mRNA sequencing reveals cellular heterogeneity and activated signaling pathways in developing cerebral cortex. Nature biotechnology, 2014. 32(10): p. 1053–1058.

22. Kolodziejczyk, A.A., et al., Single cell RNA-sequencing of pluripotent states unlocks modular transcriptional variation. Cell Stem Cell, 2015. 17(4): p. 471–485.

23. Baron, M., et al., A single-cell transcriptomic map of the human and mouse pancreas reveals inter-and intra-cell population structure. Cell systems, 2016. 3(4): p. 346-360. e4.

24. Hartigan, J.A. and M.A. Wong, Algorithm AS 136: A k-means clustering algorithm. Journal of the Royal Statistical Society. Series C (Applied Statistics), 1979. 28(1): p. 100–108.

25. Strehl, A. and J. Ghosh, Cluster ensembles--a knowledge reuse framework for combining multiple partitions. Journal of machine learning research, 2002. 3(Dec): p. 583–617.

26. Hubert, L. and P. Arabie, Comparing partitions. Journal of classification, 1985. 2(1): p. 193–218.

27. Vinh, N.X., J. Epps, and J. Bailey, Information theoretic measures for clusterings comparison: Variants, properties, normalization and correction for chance. Journal of Machine Learning Research, 2010. 11(Oct): p. 2837–2854.

28. Petropoulos, S., et al., Single-cell RNA-seq reveals lineage and X chromosome dynamics in human preimplantation embryos. Cell, 2016. 165(4): p. 1012–1026.

29. Huang, D.W., B.T. Sherman, and R.A. Lempicki, Systematic and integrative analysis of large gene lists using DAVID bioinformatics resources. Nature protocols, 2009. 4(1): p., 44.

30. Ito, K. and T. Suda, Metabolic requirements for the maintenance of self-renewing stem cells. Nature reviews. Molecular cell biology, 2014. 15(4): p., 243.

31. McCarthy D.J., et al., Scater: pre-processing, quality control, normalization and visualization of single-cell RNA-seq data in R. Bioinformatics, 2017. 33(8): p. 1179–1186.

32. Vincent, p., et al. Extracting and composing robust features with denoising autoencoders. in Proceedings of the 25th international conference on Machine learning., 2008. ACM.

33. Krizhevsky, A., I. Sutskever, and G.E. Hinton. Imagenet classification with deep convolutional neural networks. in Advances in neural information processing systems., 2012.

34. Camp, J.G., et al., Multilineage communication regulates human liver bud development., 2017.

35. Darmanis, S., et al., A survey of human brain transcriptome diversity at the single cell level. Proceedings of the National Academy of Sciences, 2015. 112(23): p. 7285–7290.

36. Deng, Q., et al., Single-cell RNA-seq reveals dynamic, random monoallelic gene expression in mammalian cells. Science, 2014. 343(6167): p. 193–196.

37. Klein, A.M., et al., Droplet barcoding for single-cell transcriptomics applied to embryonic stem cells. Cell, 2015. 161(5): p. 1187–1201.

38. Li, H., et al., Reference component analysis of single-cell transcriptomes elucidates cellular heterogeneity in human colorectal tumors. Nature Genetics, 2017. 49(5): p. 708–718.

39. Patel, A.P., et al., Single-cell RNA-seq highlights intratumoral heterogeneity in primary glioblastoma. Science, 2014. 344(6190): p. 1396–1401.

40. Usoskin, D., et al., Unbiased classification of sensory neuron types by large-scale single-cell RNA sequencing. Nature neuroscience, 2015. 18(1): p. 145–153.

41. Xin, Y., et al., RNA sequencing of single human islet cells reveals type 2 diabetes genes. Cell metabolism, 2016. 24(4): p. 608–615.

42. Zeisel, A., et al., Cell types in the mouse cortex and hippocampus revealed by single-cell RNA-seq. Science, 2015. 347(6226): p. 1138–1142.

